# The functional anatomy of nociception: effective connectivity in chronic pain and placebo responders

**DOI:** 10.1101/2024.03.10.584304

**Authors:** Sanjeev Nara, Marwan N. Baliki, Karl J. Friston, Dipanjan Ray

## Abstract

There is growing recognition of cortical involvement in nociception. The present study is motivated by predictive coding formulations of pain perception that stress the importance of top-down and bottom-up information flow in the brain. It compares forward and backward effective connectivity - estimated from resting-state fMRI - between chronic osteoarthritic patients and healthy control subjects. Additionally, it assesses differences in effective connectivity between placebo responders and non-responders and asks whether these differences can be used to predict pain perception and placebo response. To assess hierarchical processing in nociception, we defined two primary cortical regions: primary somatosensory cortex (SSC) and posterior insula (PI) (primary interoceptive cortex) and lateral frontal pole (FP1), a terminal relay station of the pain processing pathways. The directed (effective) connectivity within and between these regions were estimated using spectral dynamic causal modeling (DCM). 56 osteoarthritis patients and 18 healthy controls were included in the analysis. Within the patient group, effective connectivity was compared between placebo responders and non-responders.

In osteoarthritic patients, contra control group, forward connectivity from SSC to FP1 and from PI to FP1 was enhanced in the left hemisphere. Backward connections from FP1 to SSC were more inhibitory. Intrinsic (i.e., inhibitory recurrent or self-connectivity) of left FP1 increased. In placebo responders compared to non-responders, forward connections from bilateral SSC to PI, left SSC to FP1, left PI to left FP1 were more inhibitory. In addition, self-connections of bilateral PI and top-down connections from right FP1 to right SSC were disinhibited; whereas self-connections of right FP1 became increasingly inhibitory. We confirmed the robustness of these results in a leave-one-out cross-validation analysis of (out-of-sample) effect sizes. Overall, effective extrinsic and intrinsic effective connectivity among higher and lower cortical regions involved in pain processing emerges as a promising and quantifiable candidate marker of nociception and placebo response. The significance of these findings for clinical practice and neuroscience are discussed in relation to predictive processing accounts of placebo effects and chronic pain.

## Introduction

Pain remains a predominant reason for medical consultations, not only in most developed nations but also in developing countries.^1^ It has a profound effect on quality of life, mental health, and overall functionality. Despite its ubiquity, pain particularly in the form of chronic pain, remains enigmatic and often impervious to effective treatment.^2^ To devise more efficacious pain management strategies, it is imperative to gain a mechanistic understanding of its functional anatomy. In this study, we draw inspiration from contemporary paradigms in brain function based on hierarchical predictive coding.^3–5^ Employing dynamic causal modeling (DCM),^6,7^ we compared patients with chronic osteoarthritic pain and control participants to characterize the central (predictive) processing that underwrites nociception and the placebo response.

Neurobiological perspectives on pain have seen significant changes over the years. Traditional studies in the field adhered to the Cartesian viewpoint (pain as a bottom-up, direct-line sensory system, through which nociceptive inputs would travel to the brain, much like ringing a bell by pulling a rope).^8^ This perspective constituted the majority of animal model studies, focusing on peripheral afferents, spinal cord circuitry, their reorganization, and the molecular targets underlying chronic pain in rodent models. However, research has gradually expanded to include cerebral cortical regions,^9,10^ thanks to advances in neuroimaging techniques. Notably, functional magnetic resonance imaging (fMRI) has played a pivotal role in shedding light on the cortical areas associated with pain perception,^11^ including the somatosensory cortex, insula, anterior cingulate cortex, and prefrontal cortex.

Initially, neuroimaging research on pain perception focused on functional localization, associating specific brain regions with discrete functions related to pain processing^12^. However, this view has evolved towards a more comprehensive perspective that emphasizes distributed and synthetic processing^13^. This shift recognizes that pain perception does not exclusively rely on distinct regions but instead involves hierarchical interactions among various brain regions and networks.

Furthermore, the conventional notion of the brain as a passive receiver of incoming stimuli has been challenged by emerging theories like hierarchical predictive coding.^3–5^ These theories underscore the pivotal role of top-down cortical processing in shaping our perception of pain. Pain is a highly subjective experience, as elucidated by the definition provided by the International Association for the Study of Pain^14^ : “an unpleasant sensory and emotional ordeal linked to real or potential tissue damage, or described in the context of such harm.” Pain perception is influenced by memories, emotions, cognitive factors, and other variables, and the resulting pain experience does not necessarily correspond to linear nociceptive drive. Recent data even suggest that painful experiences can occur without a primary nociceptive input. ^15–18^ Thus, understanding the interaction between top-down processing and bottom-up sensory inputs is crucial, not only in the context of chronic pain but also in understanding its response to placebo treatment, where top-down cortical processing undoubtedly assumes a foundational role.

Motivated by the growing appreciation of the current processing in hierarchical predictive processing accounts of perceptual and active inference, our study analyses the effective connectivity - among hierarchically organized sensorimotor regions - and its characteristics in patients with chronic knee osteoarthritic pain, and their response to placebo treatment. Here, effective connectivity refers to the (directed) influence that one neural system exerts over another, either at a synaptic or population level.^19^ In contrast to data-driven approaches, such as whole-brain functional connectivity analyses, we committed to a model-based approach that allows one to test hypotheses about the functional organization of the pain network, via Bayesian model comparison.

Our aim was to identify the distinct patterns of top-down and bottom-up effective connectivity within the pain processing pathway in individuals with chronic pain, and those who respond to placebos. Hence, we chose two primary sensory cortices - and a high level (deep or terminal) node in the nociceptive pathway - as regions of interest in a minimal pain hierarchy. These regions were the primary somatosensory cortex (SSC), the posterior insula (PI) (also known as the primary interoceptive cortex), and the lateral frontal pole (FP1), respectively. Our connectivity analysis used spectral dynamic causal modeling (DCM) of resting-state fMRI data collected from chronic osteoarthritic knee pain patients and a control group. We identified connections that exhibit significant alterations in osteoarthritic patients. Additionally, in a subset of patients receiving placebo therapy, we identify distinct connections significantly associated with the placebo response. We estimated effect sizes for both analyses through leave-one-out cross-validation using parametric empirical Bayesian methods.

## MATERIALS AND METHODS

The primary aim of this research was to identify changes in extrinsic (i.e., between regions) top-down, bottom-up, and intrinsic (i.e., within region) recurrent connectivity within cortical regions that process nociceptive information in subjects suffering from chronic osteoarthritic pain, relative to a control cohort. Furthermore, we aimed to identify differences in connectivity between chronic pain patients who respond to placebo treatments from those who do not. We collected data from two independent studies, which were subject to spectral DCM. The procedural framework for our analysis is depicted in Figure 1.

**Figure 1:**
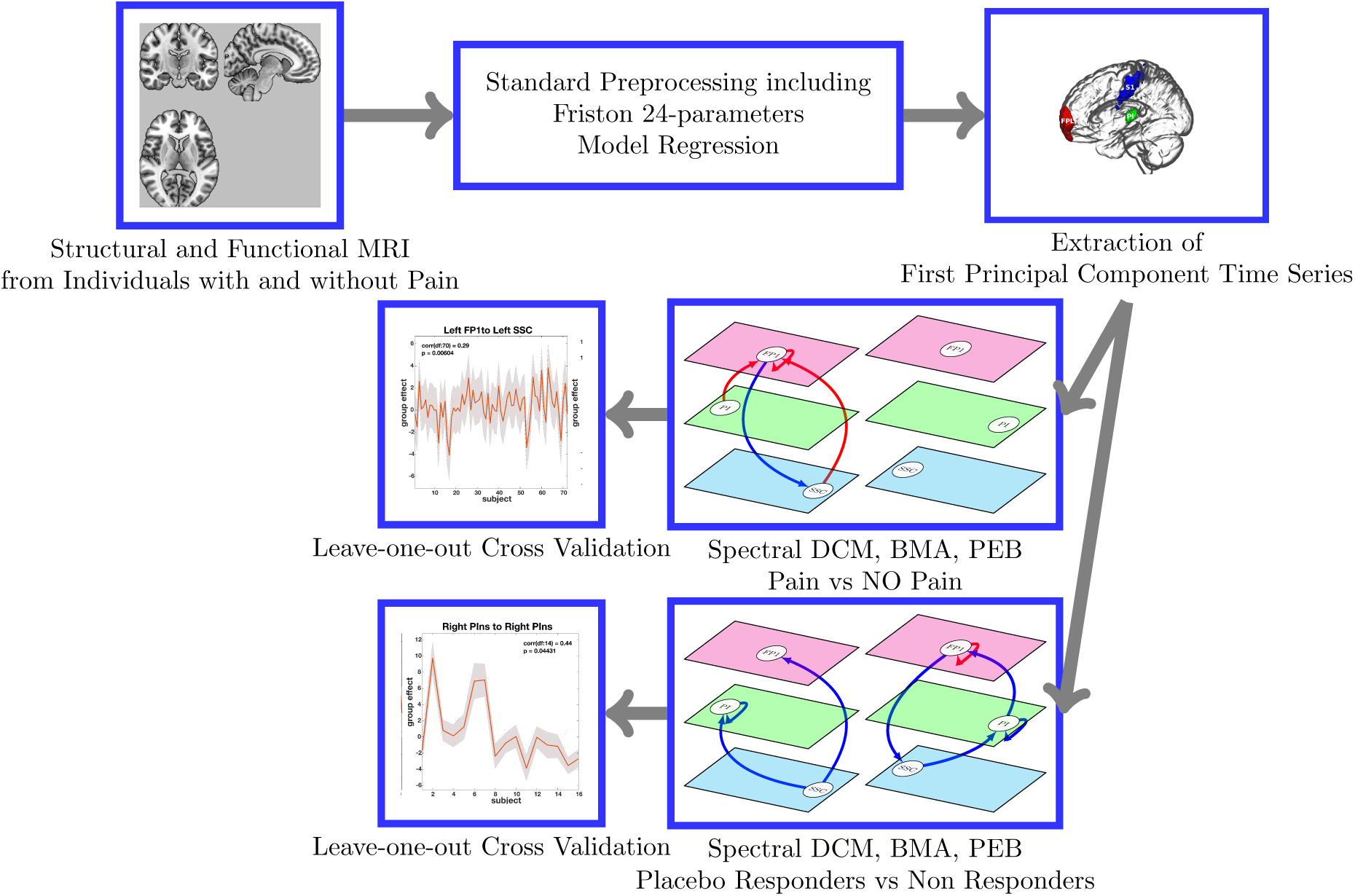
Analysis pipeline

### Study Participants

The participants in this study were drawn from two separate research projects conducted at Northwestern University, USA. The first study involved a two-week placebo-only treatment, while the second study compared three months of placebo treatment with duloxetine. A number of patients either did not complete the studies they were enrolled in, or their brain scans did not meet the quality assessment standards. As a result, the present analysis included 16 patients from study 1 and 38 patients from study 2. Additionally, 20 age-matched healthy control subjects were recruited, although data from 2 of these subjects had to be discarded due to quality concerns, leaving a final subset of 72 participants.

The patients were recruited through public advertisements and Northwestern University-affiliated clinics. All patients underwent brain scans before the commencement of their respective treatments or placebos. Each participant provided written informed consent to participate in procedures that were approved by the Northwestern University Institutional Review Board committee (STU00039556).

All osteoarthritis (OA) participants met the criteria established by the American College of Rheumatology for OA and experienced pain for at least one year. Specific inclusion and exclusion criteria were applied, including the presence of other chronic pain conditions and major depression. Participants were required to have knee pain intensity rated at least 4 out of 10 on an 11-point numerical rating scale (NRS) within 48 hours of the screening visit. A detailed list of all inclusion and exclusion criteria can be found in supplementary Document 1, with participant demographics provided in Table 1.

### Study Design

In the current study, we analyzed a subset of data from two studies. Our primary goal was to compare resting-state effective connectivity patterns among patients from both studies and control participants. Additionally, we conducted comparisons of effective connectivity between placebo responders and non-responders in Study 1. It is important to note that we refrained from merging the two studies in the later analysis due to variations in the duration of placebo use (two weeks versus three months).

For brevity, we have omitted specific details about the two studies in this paper. Readers interested in obtaining further information can refer to (T’etreault et al., 2016 and Schnitzer et al., 2018)^20,21^ for details.

**Table 1:** Demographics and clinical data.

| Groups | Study 1 |  | Study 2 |  |
| --- | --- | --- | --- | --- |
|  | Patients | Controls | Responders | Non-responders |
| Age | 57.40 ± 6.55 | 58.16 ± 6.87 | 55.87 ± 4.05 | 56.25 ± 5.57 |
| Gender | 29 F 25M | 10F 8M | 5F 3M | 3F 5M |
| Dominant Injury Side | 51 Right 5 Both | NA | 8 Right | 8 Right |
| Baseline VAS | 6.77 ± 1.41 | NA | 7.37 ± 1.35 | 6.68 ± 0.75 |
| Baseline WOMAC | 45.96 ± 15.27 | NA | 45.50 ± 18.60 | 36.25 ± 14.49 |
| % Analgesia VAS | 18.84 ± 33.35 | NA | 54.34 ± 29.45 | -7.12 ± 18.58 |
| % Analgesia WOMAC | 19.63 ± 30.85 | NA | 38.61 ± 24.60 | 7.34 ± 20.97 |

### Behavioral and Clinical Measures

Participants from both studies completed a general health questionnaire and a Visual Analog Scale (VAS) rating of their knee OA pain, on a scale of 0 to 10. Additionally, participants completed the Western Ontario and McMaster Universities Osteoarthritis Index (WOMAC), the Beck Depression Inventory (BDI), and the Pain Catastrophizing Scale (PCS). All questionnaires were administered on the day of brain scanning. Response categorization was determined initially using only the VAS measure and then validated using WOMAC scores. Analgesic and placebo responses were pre-defined individually as a minimum of a 20% reduction in VAS pain from baseline to the end of the treatment period; otherwise, subjects were classified as non-responders. This threshold was chosen based on prior research findings^22^ and a recent meta-analysis estimating the magnitude of placebo analgesia.^23^ In Study 2, to partially account for regression to the mean effects, VAS pain was measured three times over a two-week period before treatment initiation and after discontinuation of medication use, with the average score used as the indicator of pain at study entry.

### Neuroimaging Data

Figure 1 offers a schematic outlining the steps involved in the acquisition and analysis of neuroimaging data. The neuroimaging data from a subset of participants in the current study has been previously reported in another publication.^20^ However, that earlier study primarily used a data-driven approach based on correlation-based (undirected) functional connectivity analysis of whole-brain data. In contrast, the current study addresses specific hypotheses by evaluating the evidence for network models of (directed) effective connectivity among functionally characterized brain regions.

### Functional MRI data acquisition

Imaging data were acquired using a 3T Siemens Trio scanner equipped with a standard radio-frequency head coil. The structural and functional images were acquired with the following parameters: Structural MRI: Sequence: MPRAGE (T1-anatomical brain images), Field of View: 256 × 256 × 256 mm, TR/TE: 2500/3.36 ms, Flip Angle: 9°, Voxel Size: 1 × 1 × 1 mm, Slices: 160 Functional MRI: Sequence: Multi-slice T2*-weighted echo-planar images, TR/TE: 2500/30 ms, Flip Angle: 90°, Slice Thickness: 3 mm, In-plane Resolution: 64 × 64, Number of Slices: 40

### Pre-processing

The fMRI data underwent preprocessing and analytical procedures using the SPM12 v7771 toolbox (Statistical Parametric Mapping, available at http://www.fil.ion.ucl.ac.uk/spm). To ensure equilibrium in magnetization, the initial five scans were excluded. The preprocessing pipeline included slice timing correction, alignment to the mean image, motion correction, and coregistration with the participant’s T1-weighted scans. Subsequently, the images were normalized to the standard Montreal Neurological Institute (https://www.mcgill.ca) template, and resampled to 4 × 4 × 5 mm³ resolution. Motion correction involved second-degree B-Spline interpolation for estimation and fourth-degree for reslicing, while coregistration leveraged an objective function based on mutual information, and spatial normalization used fourth-degree B-Spline interpolation. Spatial smoothing was applied with a Gaussian kernel at full-width half-maximum dimensions of 4 × 4 × 10 mm³. Additional noise reduction was performed by regressing out extraneous variables, including Friston-24 head motion parameters and signals from the cerebrospinal fluid and white matter. To mitigate low-frequency drifts in the data - stemming from physiological activities and scanner-related factors - temporal filtering with a high-pass threshold of 1/128 Hz was applied.

### Selection of ROIs and extraction of time series

The areas we focused on in our study included the lateral frontal pole (FP1), the primary somatosensory cortex (SSC), and the posterior insula (PI), as shown in Figure 2. We defined these regions of interest by using predefined masks from the SPM Anatomy toolbox, as cited in reference.^24^ To prepare the data for dynamic causal modeling (DCM), we took the first principal components of the voxel time series within these masks. We then adjusted the time series for “effects of interest” (i.e., mean correcting the time series).

**Figure 2:**
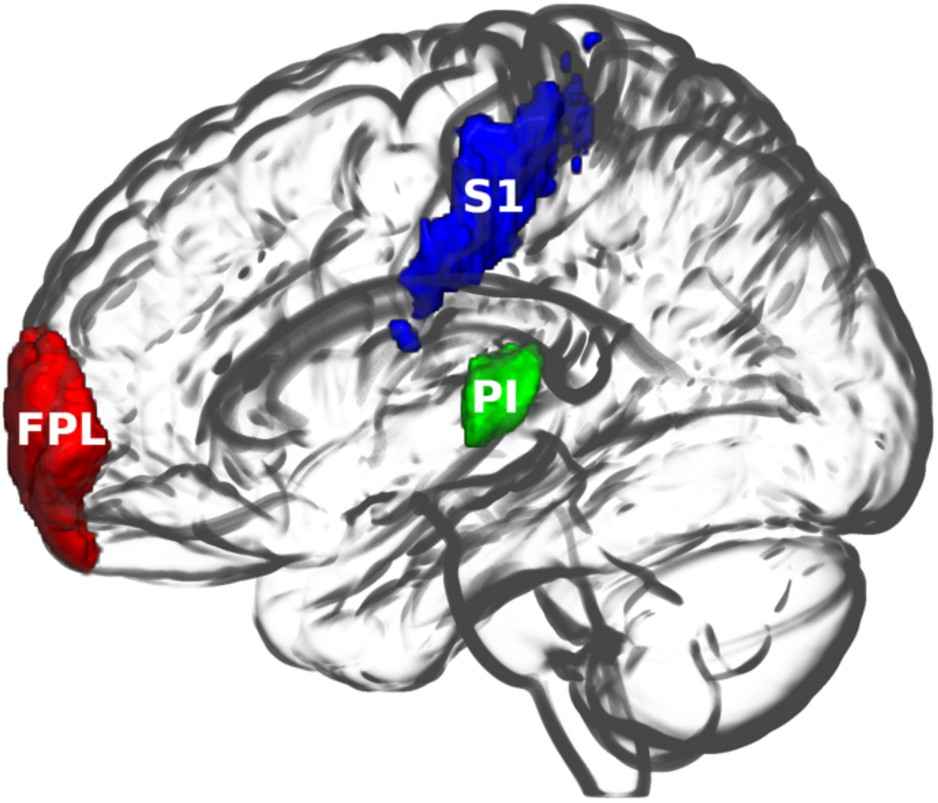
Regions of interest: FPL: lateral frontal pole, S1: primary somatosensory cortex, PI: posterior insula. The images were created using MRIcroGL (https://www.nitrc.org/projects/mricrogl/).

### Dynamic Causal Modelling and Parametric Empirical Bayes

We used spectral DCM implemented in SPM12 v7771 (http://www.fil.ion.ucl.ac.uk/spm) to estimate effective connectivity within and between brain regions in the above (minimal) pain hierarchy. Spectral DCM offers computational efficiency, when estimating effective connectivity from resting state timeseries, which are summarized in terms of their cross spectral density. Dynamic Causal Modeling^6^ represents a well-established technique for estimating the causal architecture (i.e., directed effective connectivity) generating distributed neuronal responses using observed BOLD (Blood-Oxygen-Level-Dependent) signals recorded from fMRI. This approach relies primarily on two equations. First, the neuronal state equation models the change in a neuronal activity over time, considering directed connectivity within a distributed set of regions. In the context of DCM for cross spectral density^7^, these regions are subject to endogenous fluctuations, where the requisite spectrum is estimated. Second, an empirically validated hemodynamic model describes the transformation of the neuronal state into a BOLD response.

The neural state equation can be expressed as follows:

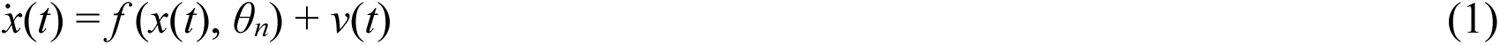

The function *f* represents the neural model in terms of neuronal dynamics, *x* represents the rate of change of neuronal states *x*, *θ_n_* signifies the unknown connectivity parameters, reflecting effective connectivity and *v*(t) accounts for a stochastic process that models endogenous neuronal fluctuations, driving the resting state. The hemodynamic model equation converts the ensuing neuronal state into a BOLD measurement:

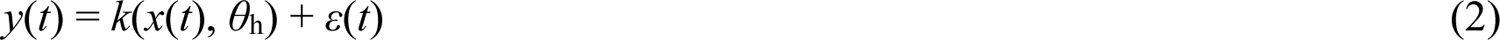

Here, the function *k* defines the biophysical mechanisms responsible for translating neuronal activity into the BOLD response, characterized by parameters *θ*_h_, and also accounts for measurement noise denoted as *ε*.

Spectral DCM^7^ provides a computationally efficient approach to invert models for resting state fMRI. It simplifies the generative model estimation process by transforming data features into the frequency domain using Fourier transforms, as opposed to using the original BOLD time series. By utilizing second-order statistics, specifically complex cross-spectra, spectral DCM overcomes the challenge of estimating time-varying fluctuations in neuronal states. Instead, it estimates their spectra, which remain time-invariant. In essence, this approach replaces the complex task of estimating hidden neuronal states with the more manageable problem of estimating their correlation functions of time or spectral densities across frequencies, including observation noise. To achieve this, a scale-free (power-law) formulation is utilized for both endogenous and error fluctuations, as outlined in Bullmore et al., 2001, ^25^ expressed as follows:

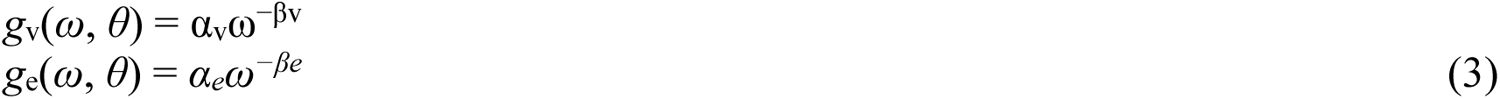

Here, the parameters *α, β* ⊂ *θ* determine the amplitudes and exponents that control the spectral density of these random effects. We employ a standard Bayesian model inversion, specifically the Variational Laplace method, to estimate the model parameters based on the observed signal, encompassing both the parameters related to the fluctuations and the effective connectivity. For a comprehensive mathematical explanation of spectral DCM, please refer to (K. J. Friston et al., 2014)^7^ and (Razi et al., 2015). ^26^

In our first-level (i.e., within subject) analysis, we estimated fully connected models for each subject within the nociceptive network in both hemispheres (right and left). Each network comprised three nodes, and we estimated both between-node (extrinsic) and within-node (intrinsic) effective connectivity. To assess the accuracy of model inversion, we examined the average percentage of variance explained by the DCM model when applied to the observed (cross-spectral) data.

For our second-level (i.e., between subject) analysis, we employed Parametric Empirical Bayes (PEB); namely, a hierarchical Bayesian model with a general linear model (GLM) of subject-specific parameters. The purpose of this PEB analysis is to estimate the effects of pain or placebo responses on effective connectivity, with subject-specific connections as random effects^27,28^.

PEB offers advantages over classical tests based upon summary statistics as it incorporates the full posterior density over parameters from each subject’s DCM, including their posterior expectations and associated uncertainties^29^. By default, the group mean serves as the first regressor in the PEB GLM. Additionally, our analysis included three more regressors: group membership (patient vs control or placebo responders vs non-responders), age, and sex. We employed Bayesian model reduction (BMR) to explore the range of potential models capable of explaining the resting state data across all subjects. BMR assesses candidate models by iteratively removing one or more connections from a full or parent model, as outlined in Friston et. al. 2016. ^28^ This process involves pruning connection parameters from the full model and evaluating the change in log model-evidence. The pruning continues until no further improvement in model evidence is observed. Finally, we addressed uncertainty over the remaining models using Bayesian Model Averaging (BMA), as described in Penny et al., 2010.^30^ BMA combines the parameters of selected models and averages them proportionally, based on their model evidence.

### Leave-one-out cross-validation Analysis

In the concluding part of our analysis, we asked whether the individual variations in effective connectivity could serve as predictors for pain perception and placebo response (in those who perceive pain). Essentially, we sought to determine if the effect size was sufficiently large to have predictive validity for a new subject. Selection criteria for connections included those that met a 95% posterior probability threshold (i.e., strong evidence) from Bayesian model reduction above. Employing a leave-one-out cross-validation method, as detailed in reference Friston et. al., 2016,^28^ we excluded one subject at a time. The predictive model, based on a parametric empirical Bayesian framework, was then applied to estimate the probability of the excluded participant’s classification (i) experiencing pain or not, and (ii) responding to placebo or not), using the previously selected connections. The accuracy of the model’s predictions was quantified by computing the Pearson’s correlation between the actual and predicted classifications of group membership.

## RESULTS

### Accuracy of DCM model estimation

The inversion of DCM models for individual participants produced excellent results in terms of accuracy (see Figure 3). Across participants, the mean variance-explained by DCM - when fitted to observed (cross spectra) data - were 80.36% (median 85.27 %) and 77.16% (median 83.68 %) for right and the left hemisphere, respectively.

**Figure 3:**
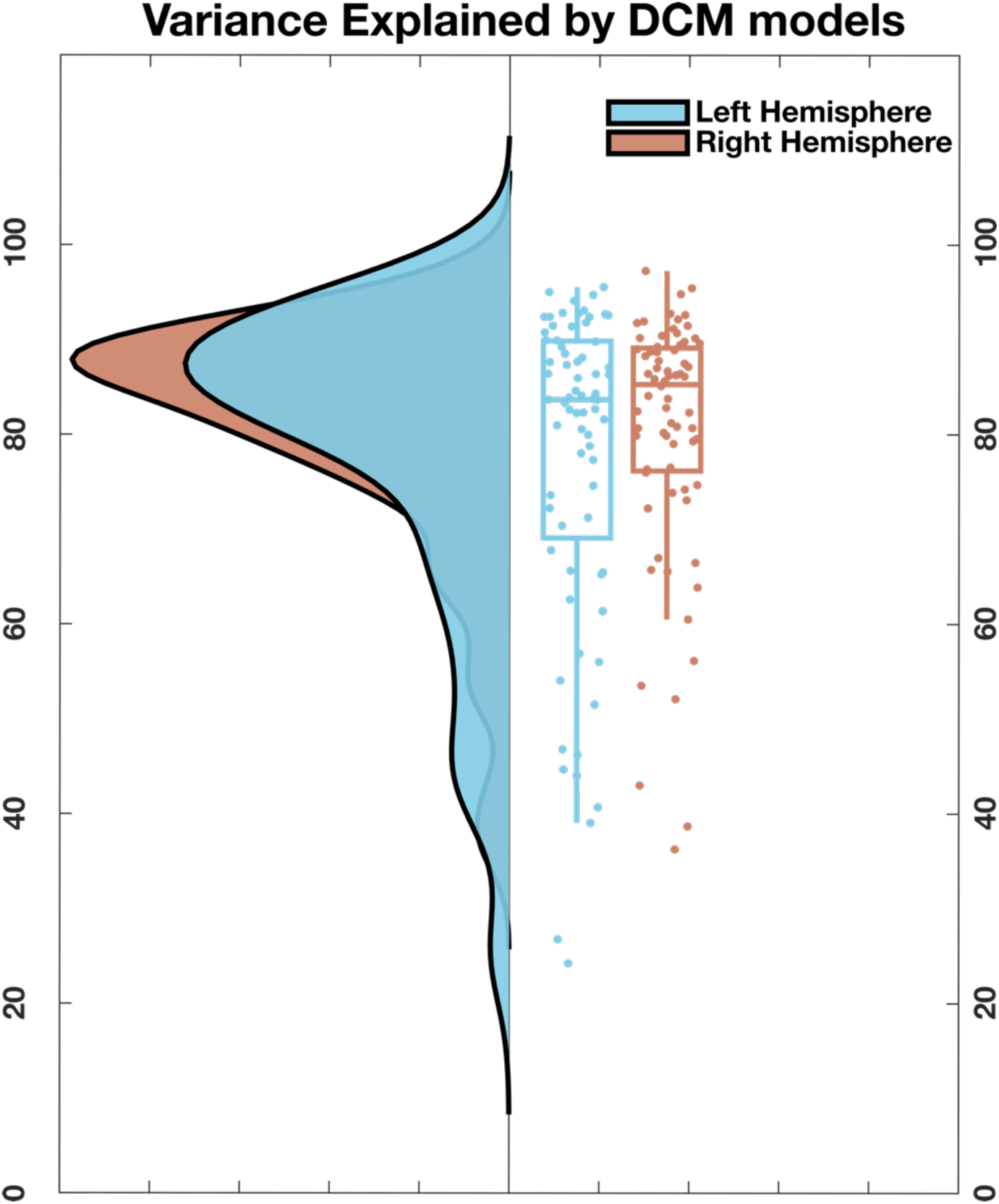
Accuracy of DCM model inversion: Average-percentage explained by our DCMs, for the target networks in both hemispheres.

### Effective connectivity

The quantitative estimates of effective connectivity for both studies are summarized in Figure 4a. These estimates of directed coupling are in units of hertz (per second) for extrinsic (between region or off-diagonal entries). In other words, they score the rate at which one region responds to the neuronal activity in another. The intrinsic (within region or diagonal entries) are log scaling estimates of recurrent self inhibition; such that a positive value denotes an increase in inhibitory intrinsic connectivity.

The mean connectivity (left panels) over all subjects was remarkably consistent over both studies and hemispheres. The regions have been arranged so that the lower diagonal entries reflect forward or bottom-up extrinsic connections; while the upper diagonal entries report backward or top-down extrinsic connections. These zero entries correspond to connections that have were considered redundant, following Bayesian model reduction. One can see that in every instance, top-down connections are either weak or excitatory, reflecting a positive modulatory influence on hierarchically lower regions. In accord with the no-strong-loops hypothesis^31,32^ - and the recurrent message passing implied by hierarchical predictive coding schemes^33,34^ - the corresponding forward connections are universally weak or inhibitory. The interpretation of these estimates of directed or effective connectivity should be in the context of the neuronal activity measured by fMRI, which can be thought of as a lumped metric of macroscopic neuronal population activity, and its excursions from steady-state dynamics. Here, these excursions can be attributed to interoceptive inference, and introspective (endogenous) activity associated with the resting state.

### Modulation of effective connectivity by chronic pain

The pattern of modulation of effective connectivity in chronic pain patients, in relation to controls, is illustrated schematically in Figure 4b, based on the quantitative estimates (in hertz) provided in Figure 4a. When comparing osteoarthritic patients to the control group, our analysis revealed a marked increase in forward connections from the left primary somatosensory cortex (SSC) towards the left frontal pole (FPL), as well as from the left posterior insula (PI) towards the left FPL. These differences constitute a 50% decrease in inhibitory influences; namely, a selective disinhibition of forward connectivity.

Conversely, the backward connections originating from the left FPL to the left SSC exhibited a decrease in weak backward excitatory influences, while self-connections within the left FPL became increasingly inhibitory (by about 9%). Notably, no discernible changes in effective connectivity were among or within regions in the right hemisphere. In summary, there was a left lateralized increase in forward connectivity best characterized as a disinhibitory effect in the pain group, with weaker decreases in backward connectivity.

**Figure 4.**
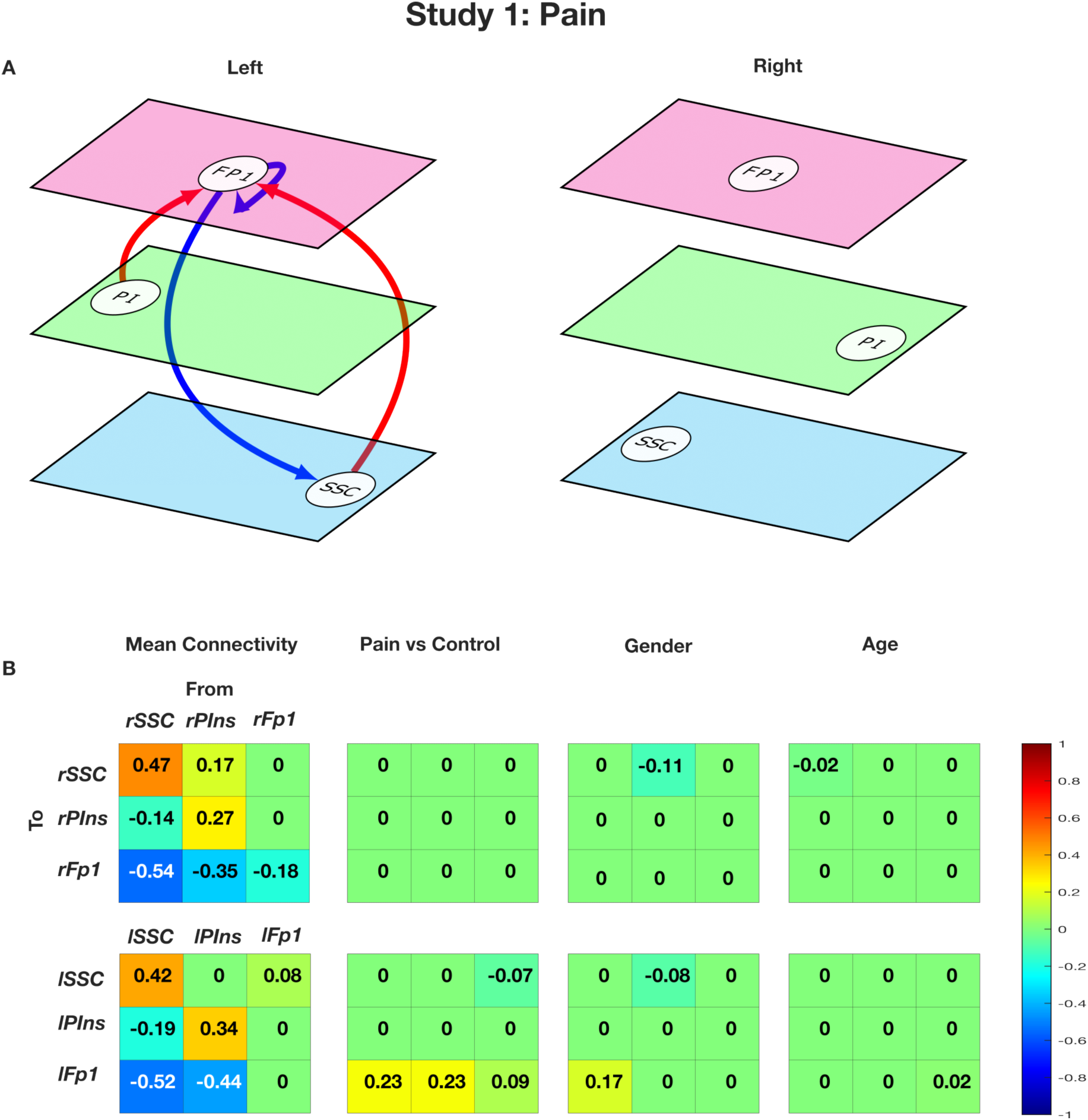
A): Modulation of effective connectivity in the patient population compared to the control group (left and right hemispheres) Arrow colors code the direction of connectivity changes relative to the group mean: red, increased; blue, decreased. For all subfigures, line thickness is kept constant and does not code for the effect size. For the exact values of the estimated connectivity parameters see Appendix A. Nodes are placed in different planes to denote relative position of different nodes in cortical hierarchy. FP1: lateral frontal pole, PI: posterior insula, SSC: primary somatosensory cortex 4B): Estimated connectivity parameters in study 1.

### Modulation of effective connectivity in placebo responders vs non-responders

Figure 5a presents a schematic reporting the relative changes in effective connectivity among individuals who exhibited a significant response to the placebo intervention. In this comparison, the findings were remarkably consistent across both hemispheres with, universally, a decrease in extrinsic connectivity, predominantly in forward connections. Specifically, in our comparison of placebo responders and non-responders, responders exhibited a shift towards an increased inhibitory influence in several key connections, including forward connections from the bilateral primary somatosensory cortex (SSC) to the posterior insula (PI), left SSC to frontal pole (FPL), left PI to FPL. This was complemented by a decrease in inhibitory self-connections within the bilateral PI. Furthermore, our analysis revealed that in responders, backward connections from the right FPL to SSC exhibited increased inhibitory effects, while self-connections within the right FPL became more inhibitory. In short, people who respond to placebo have a relative reduction in forward connectivity and increased intrinsic excitability (i.e., disinhibition) of the bilateral insular.

**Figure 5.**
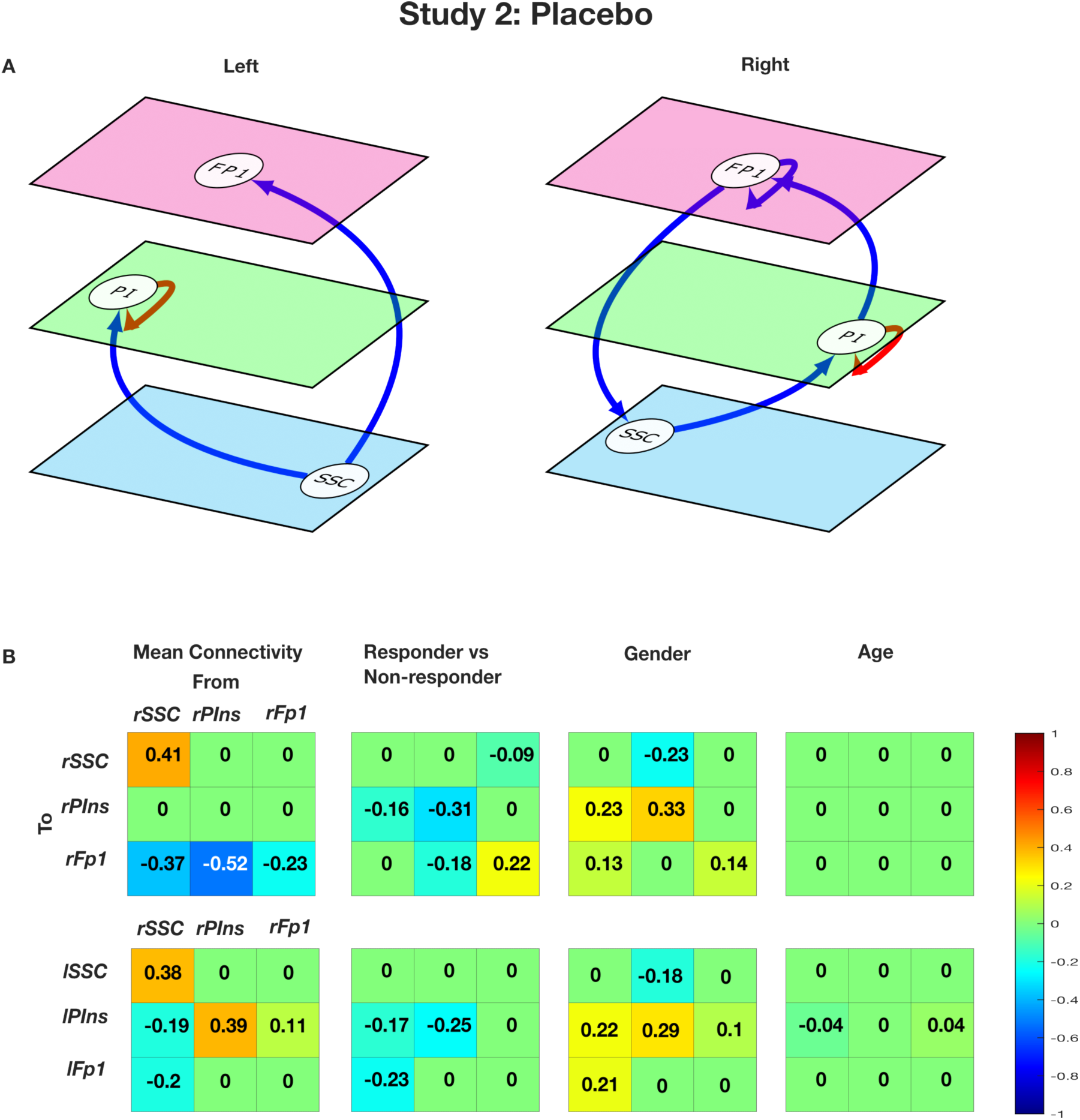
A): Modulation of effective connectivity in placebo responders vs non-responders (left and right hemispheres) Arrow colors code the direction of connectivity changes relative to the group mean: red, increased; blue, decreased. For all subfigures, line thickness is kept constant and does not code for the effect size. For the exact values of the estimated connectivity parameters see Appendix A. Nodes are placed in different planes to denote relative position of different nodes in cortical hierarchy. FP1: lateral frontal pole, PI: posterior insula, SSC: primary somatosensory cortex. 5B): Estimated connectivity parameters in study 2.

### Cross validation

In a leave-one-out cross-validation - among all connections showing significant association with chronic pain perception - directed connectivity between left FP1 to left SSC was found to predict group membership (osteoarthritis patient or control) at a significant level of α = 0.05 (see Table 2). Similarly, in another leave-one-out cross validation analysis, intrinsic connectivity within right Posterior Insula (rPIns) was found to predict placebo response among pain perceivers at the same significant level (see Table 3). This suggests a nontrivial out-of-sample effect size that is conserved over subjects.

**Table 2:** Pain Perception: Leave-one-out cross validation.

| Connections | Correlation | p-value |
| --- | --- | --- |
| ISSC → FPI | 0.15 | 0.097 |
| IPIns → IFPI | -0.18 | 0.93 |
| **IFPI → ISSC** | **0.29** | **0.006** |
| IFPI → IFP1 | 0.14 | 0.11 |

**Table 3:**
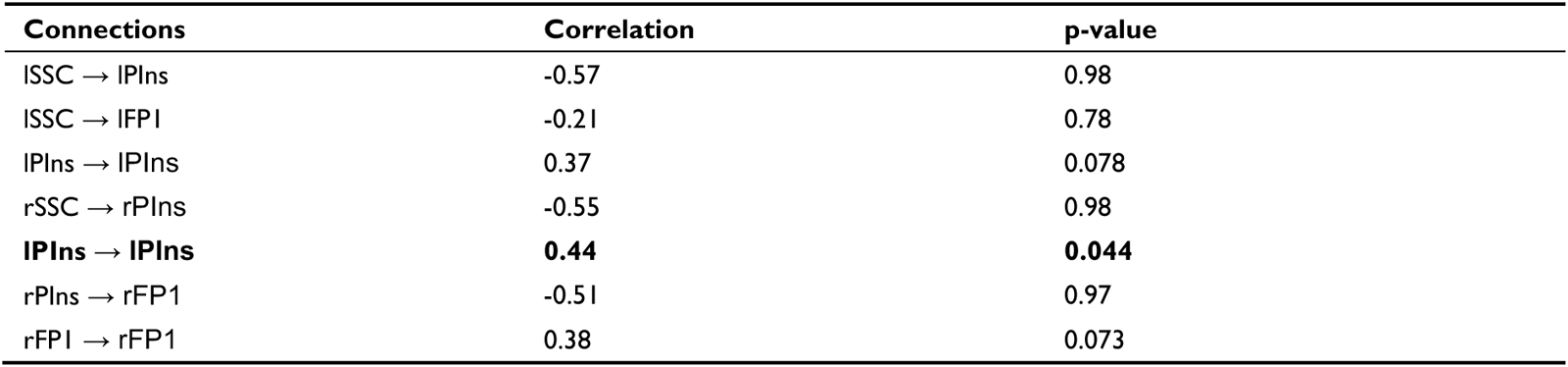
Placebo Response: Leave-one-out cross validation.

## DISCUSSION

The most striking result of our study includes the following findings: recurrent effective connectivity within the lateral frontal pole becomes more inhibitory, while backward effective connectivity (from higher to lower cortical regions) decreases in both pain perceivers (as opposed to non-perceivers) and placebo responders (compared to non-responders). However, opposite changes are observed in forward connections, where nociception is associated with more excitatory (disinhibited) connections, while placebo responses evince more inhibitory forward connections. The changes in effective connectivity among pain perceivers were only observed in the left hemisphere. In leave-one-out cross-validation analyses, we found that the top-down connection from the left FP1 to the left SSC exhibited a sufficiently large (out-of-sample) effect size to predict whether an individual is experiencing knee pain or not. In a similar analysis of placebo response, we observed that the self-connection of the right insular demonstrated a sufficiently large effect size to predict placebo responses.

There are a few neuroimaging studies that have explored changes in functional and effective connectivity within nociceptive brain regions as potential biomarkers for chronic pain and placebo response.^35–40^ However, our research takes a unique approach, driven by a novel perspective on the neural mechanisms underlying sensory perception. There is a growing consensus that perception is not simply a passive process of progressively abstracting sensory input in a “bottom-up” manner. Instead, it involves both forward and backward information flow between brain regions organized in a hierarchical fashion, which plays a pivotal role in shaping perception. This concept forms the foundation of much of the current thinking about the functional architecture of the brain. One prominent theory in this realm is predictive coding,^3–5^ which has also been extended to the domain of motor function.^19,41^ The implicit exchange of neuronal messages within cortical hierarchies motivated are characterization of effective connectivity among pain processing regions organized hierarchically, and its potential relationship with pain perception and placebo response.

The modulation of effective connectivity in osteoarthritic patients and in placebo responders have some commonalities (for example similar changes in backward and recurrent connections in the highest node, i.e., lateral pole) and some divergence (for example, opposite changes in forward connections). These changes are consistent with predictive coding accounts of pain perception and placebo response. For example, our study found that, in patients with osteoarthritic pain, top-down connections become more inhibitory and bottom-up connections more excitatory in the network involving interoceptive and somatosensory regions. These differences are consistent with the role of top-down predictions explaining away prediction errors at lower levels, as proposed by the predictive coding framework.^5,42^

In particular, they are what would be predicted in terms of hierarchical predictive coding in which precision weighted prediction errors are passed forward to deeper levels of the interoceptive hierarchy to update or revise representations at higher levels.^43–46^ An increase in forward connectivity can be read as an increase in the sensitivity of higher levels to ascending prediction errors. This corresponds to an increase in the precision of ascending prediction errors in people who experience pain or, conversely, an effective decrease in the precision of nociceptive prediction errors in people who show a placebo response. We will return to the important notion of precision and its neurophysiological correlates below.

Although long-range connections in the brain are excitatory (i.e., glutamatergic), predictive coding proposes that backward connections may preferentially target inhibitory interneurons in superficial and deep layers to evince an overall decrease in neuronal message passing.^5,42,47^ In predictive coding, this is often read as ‘explaining away’ prediction errors at lower levels in sensory cortical hierarchies so that only those incoming stimuli that deviate from prediction (i.e., prediction errors) ascend the hierarchy to revise presentations at higher levels.^42,47–50^ However, top-down predictions also predict the reliability or precision of prediction errors at lower levels, leading to a disinhibitory modulation of lower-level activity in populations encoding prediction errors: sometimes discussed in terms of attention^51^ or retrieval^52^. One can associate this effect of top-down modulation with the group mean positive modulatory effects reported above. Thus, effective connectivity in chronic pain patients appear to reflect an enhanced pain processing within the nociceptive pathway with an increase in forward connectivity corresponding to an increase in the effects of ascending or bottom-up prediction error signaling.

An intriguing finding in connectivity changes in chronic pain patients - that warrants further comment - is that differences in connectivity are restricted within the left hemisphere, without any notable changes observed in the right hemisphere. However, note that in nearly all of the patients analyzed in the present study, osteoarthritis was localized to the right knee, with only a handful experiencing bilateral knee involvement. The observation aligns with the anatomy of second-order pain neurons crossing over to the opposite side of the spinal cord and thus affording a potential explanation for left lateralized changes in connectivity.

Current formulations of nociception - and in particular, the placebo effect-rest upon predictive coding and active inference accounts of hierarchical processing within the somatosensory and interoceptive hierarchy.^45,46,53^ In particular, there is a focus on nuancing the perception of pain by adjusting the confidence or precision associated with the implicit (Bayesian) belief updating.^53–57^

In brief, it may be the case that placebo effects can be attributed to a decrease in sensory precision of the kind associated with sensory attenuation.^53,58,59^ Or, equivalently, an increase in the (subpersonal) confidence or precision afforded prior beliefs induced by the administration of placebos. Both or either of these changes in precision will move posterior beliefs towards a prior expectation that “I am not in pain because I have taken an analgesic”. In terms of predictive coding, this would correspond to an increased gain or precision weighting of ascending somatosensory and nociceptive prediction errors, relative to the precision of prior beliefs deeper in the interoceptive hierarchy (e.g., anterior insular and other prefrontal regions). From a psychological perspective, increases and decreases in the likelihood or sensory precision can be associated with selective attention or sensory attenuation, respectively. Physiologically, this kind of top-down precision weighting is thought to be mediated by selective changes in the postsynaptic sensitivity of certain neuronal populations: e.g., superficial pyramidal cells encoding prediction errors.^33,44^ It is precisely (sic) this modulation of synaptic excitability that is measured by effective connectivity and evident in our DCM results.

In short, the changes in effective connectivity are consistent with a predictive coding formulation, in the following sense. Increased bottom-up prediction error corresponds to heightened pain perception, simply because the ascending prediction errors have been afforded more precision and therefore have more influence on belief updating processes at higher levels of the hierarchy. Similarly, reduced bottom-up prediction error signaling - combined with increased top-down predictions are characteristic of placebo responders - suggesting that an increase in the precision or synaptic gain at higher hierarchical levels mitigates the accumulation of weak evidence (i.e., imprecise nociceptive prediction errors) for the high level belief: “I am in pain”.

The model comparison discussed above furnishes clear evidence for changes in a number of connections that underwrite nociception and placebo response. One might ask whether these changes can be used diagnostically in individual patients. In other words, are the underlying effect sizes sufficiently large to predict whether somebody is a patient or a control? Or anticipate the placebo response among patients? This question goes beyond whether there is evidence for an association and addresses the utility of connectivity phenotyping for precision medicine. One can estimate out of sample effect sizes using cross validation under parametric empirical Bayesian schemes.^28^ In this analysis, we withheld a particular participant and asked whether one could have predicted the group membership, given the effective connectivity estimates from that subject. In the current analysis, every connection showed a significant out-of-sample correlation with group membership for patient vs control and placebo responders vs non responders analysis. This suggests that a nontrivial amount of variance in the group membership could be explained by effective connectivity.

A note on our choice of network nodes: as we were primarily interested in quantifying top-down or backward and bottom-up or forward connectivity in the cortical hierarchy, we selected two primary sensory cortices from brain regions sensitive to pain perception and one of the highest regions in pain pathway. Thus, bilateral primary somatosensory cortices and posterior insula were selected as lower nodes. It should be pointed out here that posterior insula is widely considered as the primary interoceptive cortex.^60–62^ As higher regions we chose the right and left lateral frontal pole. Several tracing, lesion, and physiological studies suggest that visual, auditory, and somatosensory processing pathways converge at different regions of VLPFC^63,64^ and DLPFC. ^65,66^ We therefore chose the lateral frontal pole as representative of a higher node.^67^

Empirical studies^68,69^ support a posterior to anterior sensory representational hierarchy in the prefrontal cortex and place the lateral frontal pole one level higher than both DL and VL PFC in the cortical hierarchy. Involvement of lateral frontal pole in pain perception is well established in several neuroimaging and magnetic stimulation studies.^70–72^ Thus, we examined changes in the overall top-down and bottom-up effective connectivity - with pain perception and placebo response - by selecting nodes at both the highest and lowest levels of the cortical hierarchy in the pain processing pathway.

The findings from the current study should be interpreted in light of certain limitations. Firstly, our study focused on patients with knee osteoarthritis, a well-known cause of chronic pain. To ascertain whether the observed changes in connectivity represents a general pattern associated with chronic pain or a specific pattern linked to knee osteoarthritis pain, it is essential to replicate this analysis in other chronic pain conditions. Secondly, when considering our connectivity analysis, it is crucial to acknowledge the potential presence of confounding factors beyond the age and sex of the participants. For instance, depression and anxiety frequently co-occur with chronic pain and may influence top-down effective connectivity in the brain. While none of our participants reported a diagnosis of major psychiatric conditions, we did not specifically rule out, or control for, the presence of subclinical depression or anxiety.

The findings from this study hold considerable promise for practical applications. Future research could be aimed at assessing the efficacy of therapeutic interventions, encompassing various pharmacological and non-pharmacological treatments, in reversing the alterations in cortical effective connectivity and pain perception. An intriguing avenue to explore involves the use of emerging noninvasive brain stimulation techniques, such as Transcranial Magnetic Stimulation (TMS). Recent studies have demonstrated the ability of TMS to modulate cortico-cortical connectivity within specific neural circuits.^73–75^ By applying these techniques to target specific brain regions within the pain processing pathway, we can investigate their impact on nociception using state-of-the-art methodologies that are currently available.

In conclusion, our findings advance our mechanistic understanding of the development and persistence of chronic pain and the placebo response. Building upon emerging theoretical frameworks of brain function such as predictive coding, our current study highlights changes in top-down, bottom-up, and intrinsic effective connectivity in pain processing pathway as potential neural markers of nociception and the placebo response. Furthermore, it confirms the generalizability and predictive reliability of this novel marker, potentially opening up new avenues for research into the neural foundations of pain and potential therapeutic interventions.

## DATA AND CODE AVAILABILITY

Our analysis code is available on GitHub (https://github.com/dipanjan-neuroscience/pain_placebo). Imaging data are available on OpenNeuro platform (https://openneuro.org/datasets/ds000208/versions/1.0.1)

## ACKNOWLEDGEMENTS

S.N was supported by Research Cluster “The Adaptive Mind”, funded by the Excellence Program of the Hessian Ministry of Higher Education, Science, Research and the Arts. K.J.F. was supported by funding for the Wellcome Centre for Human Neuroimaging (Ref: 205103/Z/16/Z). DR’s research was made possible through the financial support provided by Ashoka university. For the purpose of Open Access, the authors have applied a CC BY public copyright license to this manuscript.

## AUTHOR CONTRIBUTIONS

S.N., and D.R. conceived the present project. M.B. was responsible for overseeing the initial data collection. D.R., and S.N. performed data analysis. K.J.F, and D.R supervised the project. S.N, and D.R., wrote the manuscript. D.R., M.B. and K.J.F edited the manuscript.

## CONFLICT OF INTEREST

The authors report no biomedical financial interests or potential conflicts of interest.

## Supplementary information 1

Full inclusion and exclusion list for both the studies was as following:

**1. Inclusion Criteria:**

- Age: 45-80 years
- ACR criteria for OA including Kellgren-Lawrence radiographic OA grades II-IV
- VAS pain score > 5/10 within 48 hrs. of the phone screen and visit 1 (Screening)
- Knee OA for a minimum of 12 months
- Need for daily pain medication to manage symptoms of OA

**2. Exclusion Criteria:**

- Currently taking MAO inhibitors or any centrally acting drug for analgesia, depression
- Narrow angle glaucoma
- Uncontrolled hypertension
- Co-existing inflammatory arthritis, fibromyalgia or other chronic pain state.
- If a female, pregnant, trying to become pregnant, or lactating
- Major depressive disorder
- Substantial alcohol use or history of significant liver disease
- Use of MAO inhibitors, triptans, serotonin precursors (tryptophan)
- Use of potent CYP1A2 inhibitors, Thioridazine, and anti-depressants
- Diabetes, type 1 or type 2
- Condition in which the Investigator believes would interfere with the subject’s ability to comply with study instructions, or might confound the interpretation of the study results or put the subject at undue risk

## Notes

https://openneuro.org/datasets/ds000208/versions/1.0.1

### Competing Interest Statement

The authors have declared no competing interest.

### Summary of Updates

Changed the formatting of title and linked orcid IDs.

